# Genomic and phenotypic characterization of *Pseudomonas hygromyciniae*, a novel bacterial species discovered from a commercially purchased antibiotic

**DOI:** 10.1101/2021.10.06.463447

**Authors:** Timothy L. Turner, Sumitra D. Mitra, Travis J. Kochan, Nathan B. Pincus, Marine Lebrun-Corbin, Bettina Cheung, Samuel W. Gatesy, Tania Afzal, Egon A. Ozer, Alan R. Hauser

## Abstract

A purchased lot of the antibiotic hygromycin B was found to be contaminated with a novel bacterial species, which we designate *Pseudomonas hygromyciniae*. Characteristics of *P. hygromyciniae* include its ability to use a variety of compounds as carbon sources, its pathogenicity towards lettuce and *Galleria mellonella*, and its ability to inhibit the growth of an *E. coli* strain. *P. hygromyciniae* is unlikely to be a human pathogen, as it did not survive at 37 °C and was not cytotoxic towards a mammalian cell line. The *P. hygromyciniae* strain harbors a novel 250 kb megaplasmid which confers resistance to hygromycin B and contains numerous other genes predicted to encode replication and conjugation machinery. These findings indicate that commercially manufactured antibiotics represent another extreme environment that may support the growth of novel bacterial species.

**IMPORTANCE:** Microbial ecologists have surveyed numerous natural and manmade environments in search of new microbial species. In some instances, these microbes are discovered in harsh conditions, such as deep-sea vents, and their discovery leads to better understanding of how microbes adapt to their environment. Here, we have discovered a new species of bacteria from an extreme manmade environment: a lyophilized, commercially available bottle of the antibiotic hygromycin B.

## INTRODUCTION

To identify novel bacterial species, microbiologists have examined a wide range of extreme niches. The basis for these investigations is that physical and biological stresses may select for bacterial species not found in more conventional environments. This strategy has been successful in a number of harsh niches such as hot springs, deep ocean trenches, and hypersalinic brine pools^1^. Continued efforts to discover novel bacterial species are important, as they can lead to the identification of novel enzymes^2^ and biomolecules^3^, and can inform our understanding of bacterial evolution.

In clinical environments, antibiotics place a strong selective pressure on bacteria. As a result, the commercial release of a new antibiotic is often followed by the emergence of infections caused by bacteria resistant to that antibiotic^4^. However, antibiotics are also found in high concentrations in some non-clinical facilities, such as sewage and certain industrial plants where antibiotics are produced. In particular, antibiotic-manufacturing facilities represent a different kind of inhospitable niche in which the harsh condition of high antibiotic concentrations provides a strong selective pressure. It is conceivable that these niches could be exploited to identify novel microbes.

Here, we describe the serendipitous discovery of a novel gram-negative bacterial species from a purchased vial of antibiotic. The new species, which we designate *Pseudomonas hygromyciniae*, was named based on its discovery from lyophilized hygromycin B, an aminoglycoside antibiotic.

## RESULTS AND DISCUSSION

While preparing selective agar for a cloning experiment, a sealed bottle of purchased lyophilized hygromycin B was opened and a portion of the powder aliquoted into sterile lysogeny broth (LB), which was then stored at 4 °C. When the LB containing hygromycin B was retrieved after several days, it was noted to contain visible microbial growth. Distinct and uniform colonies were observed when the medium was streaked onto LB agar plates with or without 500 μg/mL filter-sterilized hygromycin B. A single colony was picked from a hygromycin plate and regrown in sterile LB medium containing 500 μg/mL filter-sterilized hygromycin B, and a frozen stock was made for use in all subsequent studies. The colonies that grew on the LB agar plates were cream-colored and of a single morphology (**Supplemental Fig. 1a**) and had a pungent odor. These colonies fluoresced under a 302 nm blacklight (**Supplemental Fig. 1b**). The organism grew into visible colonies on plates incubated for 24-48 h at 4 °C, 22 °C, and 30 °C but not at 37 °C; estimated growth rate based on colony size was highest between 22 °C and 30 °C. Microscopy indicated that the organism was a gram-negative rod-shaped bacterium of approximately 1.5 μm in length and 0.5 μm in width (**Supplemental Fig. 1c**). Pending further characterization, the bacterium was referred to as strain SDM007.

Genomic characterization of SDM007 was undertaken with Illumina (short-read) and Nanopore (long-read) sequencing. Sequencing reads were assembled *de novo* to yield a complete, closed genome. The resulting sequence consisted of a 6,037,020 bp circular chromosome with G+C content of 60.2% and a 249,868 bp circular megaplasmid with a G+C content of 55.3%. A BLAST analysis of the SDM007 genome against the NCBI 16s rRNA gene sequence database for bacteria and archaea indicated that it has a unique 16s rRNA sequence most closely related to those of *Pseudomonas gessardii* (99.67% identity) and *Pseudomonas libanensis* (99.67% identity) (**Supplemental Information 1**). *In silico* DNA-DNA hybridization (DDH)^5^ was performed using the Type Strain Genome Server (TYGS)^6^ to compare SDM007 to the type strains in TYGS database. A *d_4_* value of ≤ 70% to the next nearest genome has been considered one potential indicator of a distinct species^7^. The bacteria with the highest predicted DDH to SDM007 were *Pseudomonas proteolytica* strain LMG 22710 (*d*_4_ of 60.0%), *P. gessardii* strain DSM 17151 (*d*_4_ values of 45.5%), and *Pseudomonas brenneri* strain DSM 15294 (*d*_4_ values of 45.1%). An average nucleotide identity (ANI) difference of ≤ 95% has also been used to distinguish species^7^. An approximate ANI of SDM007 with 9,108 publicly-available *Pseudomonas* spp. genomes was calculated using a previously published nucleotide similarity script^8^. The three most similar genomes from this analysis were then compared to SDM007 using the JSpeciesWS ANI-calculating software^9^. *Pseudomonas* sp. strain LG1D9_6333, *P. brenneri* BlGb0273_9434, and *P. proteolytica* BS2985_6818 had 94.5%, 94.6%, and 94.2% ANI, respectively, to SDM007. Next, a phylogenetic tree was generated from SNP loci in the core genomes (defined as sequences occurring in at least 95% of genomes) of SDM007 and the 23 most closely related *Pseudomonas* genomes with an approximate ANI ≥90%. *P. fluorescens* reference strain SBW25 was included as an outgroup to root the tree. These results illustrated the close relationship between SDM007 and *P. brenneri, P. proteolytica*, and *P. fluorescens* (**Fig. 1**). Together, the 16s ribosomal RNA gene sequencing, DDH, and ANI results indicate that SDM007 is a novel species in the *P. fluorescens* group and is closely related to *P. brenneri, P. proteolytica*, and *P. gessardii.* Based upon the source from which this bacterium was discovered, we propose that it be named *Pseudomonas hygromyciniae*.

**Figure 1.**
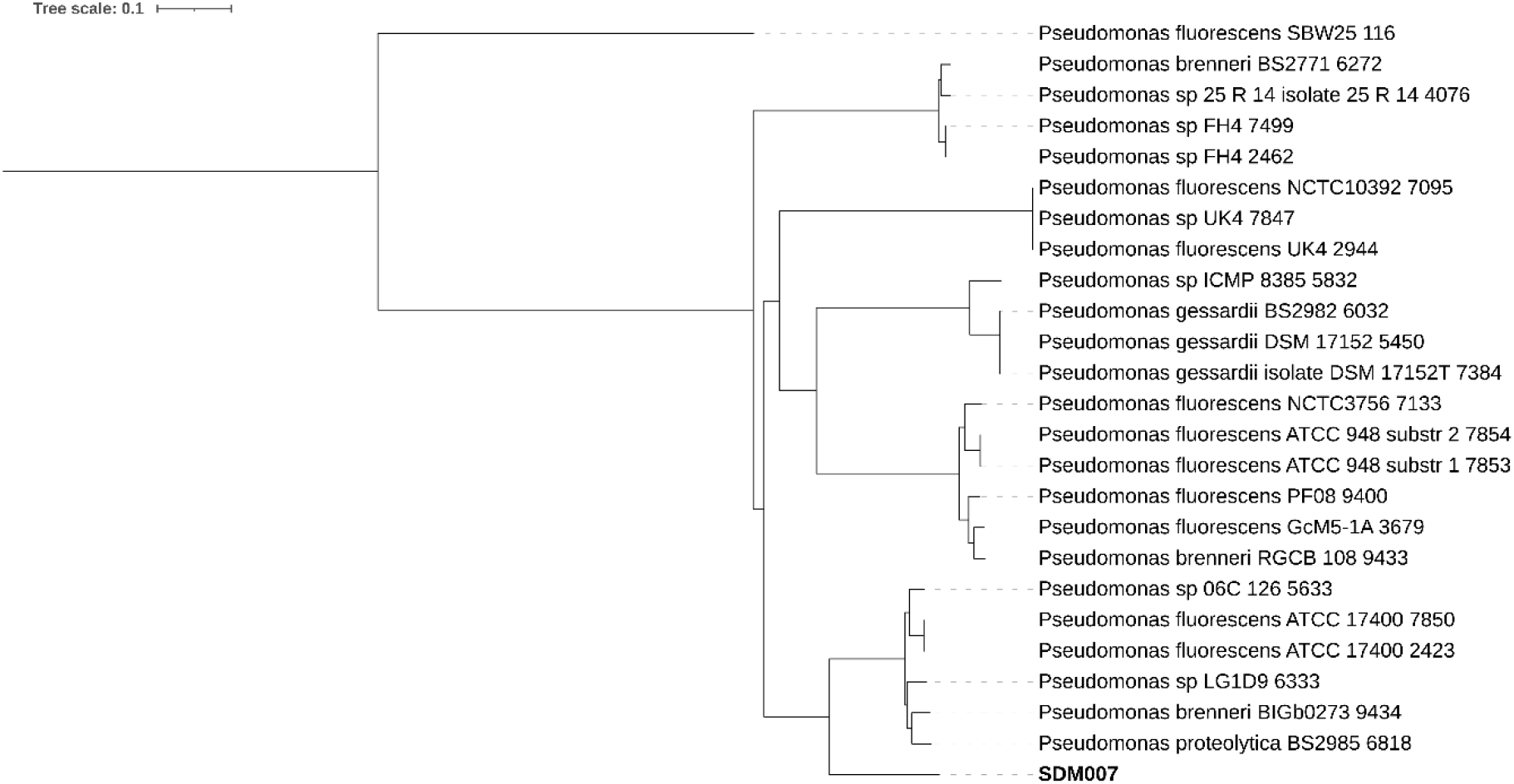
Phylogeny of SDM007 and closely related bacteria. A parsimony-based phylogenetic tree was generated based on SNP loci occurring in at least 95% of the analyzed genomes. *Pseudomonas fluorescens* reference strain SBW25 was included as an outlier to root the tree. Strain names are given as listed in the Pseudomonas Genome Database^25^ and do not necessarily represent current species designations.

To better understand how *P. hygromyciniae* may differ from other *Pseudomonas* spp., a variety of phenotypic assays were conducted to compare this bacterium to *Pseudomonas aeruginosa* and *P. fluorescens*. Optical density measurements at 600 nm (OD_600_) suggested that *P. hygromyciniae* grew at a rate comparable to the *P. fluorescens* strain ATCC 17569 at 4 °C, while *P. aeruginosa* strain PAO1 did not show appreciable growth at this temperature (**Fig. 2a**). At 30 °C, *P. hygromyciniae* and *P. aeruginosa* PAO1 showed similar growth kinetics, and *P. fluorescens* ATCC 17569 grew slightly faster (**Fig. 2b**). Notably, *P. hygromyciniae* failed to show detectable growth at 37 °C after 24 hours, unlike *P. fluorescens* ATCC 17569 and *P. aeruginosa* PAO1 (**Fig. 2c**). Growth rates were then confirmed by enumeration of CFU from streaked aliquots of liquid cultures. Although *P. hygromyciniae* grew relatively well at 4 °C (**Fig. 2d**) and had comparable growth to the control strains at 30 °C (**Fig. 2e**), it rapidly died at 37 °C, with a decrease of approximately 1 log of viable CFU after 12 hours of incubation (**Fig. 2f**). Unsurprisingly, *P. hygromyciniae* was resistant to hygromycin B, as it grew from an initial OD_600_ of ~0.1 to over 1 within 20 hours at 30 °C in LB containing 500 μg/mL of this antibiotic (**Fig. 2g**). No statistically significant difference in OD_600_ was observed at the 24 h timepoint for SDM007 grown in LB medium supplemented with 500 μg/mL vs. 5 mg/mL hygromycin B, although the growth rate was slower over the first 8 hours in medium containing the higher concentration (**Fig. 2h**).

**Figure 2.**
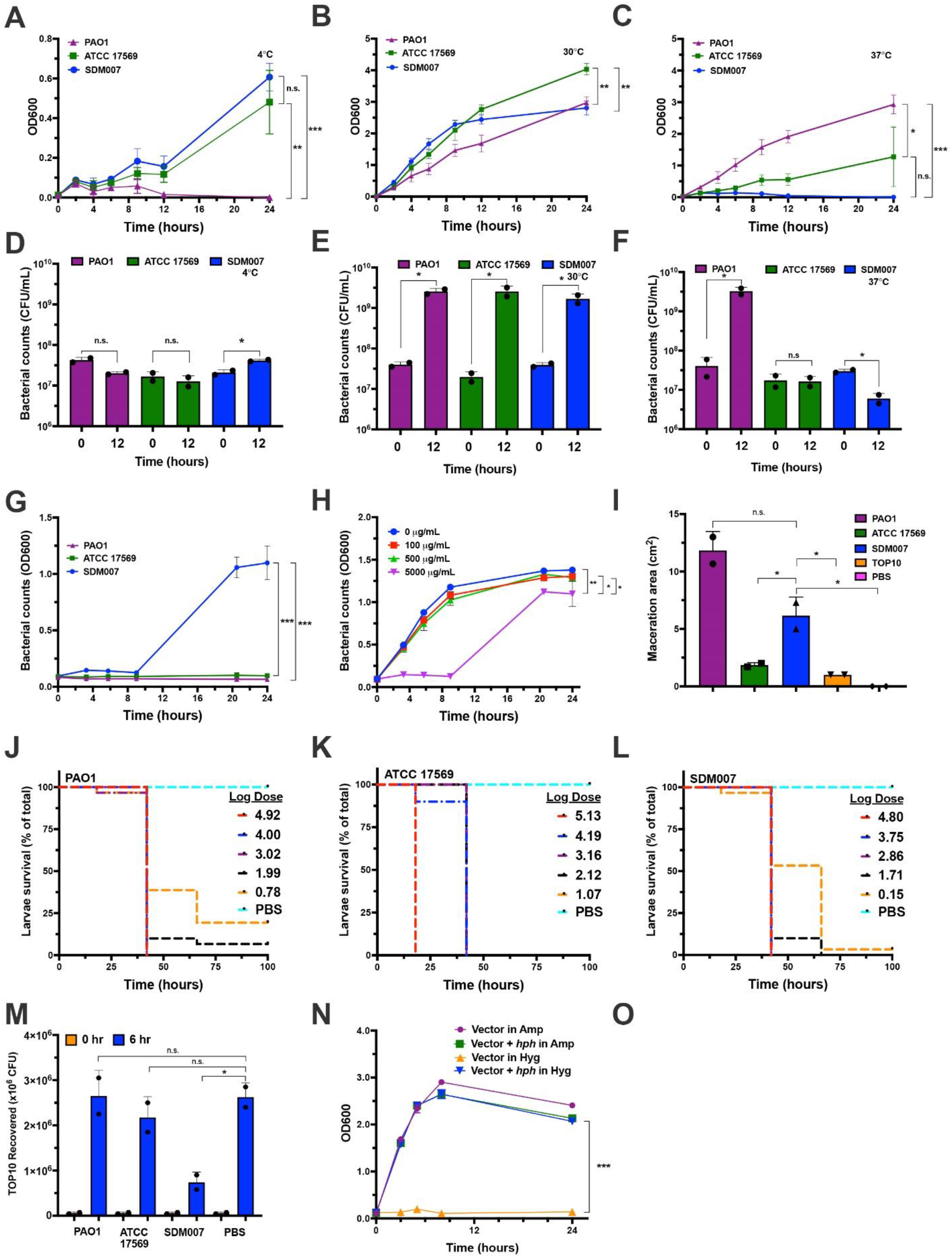
Phenotypic characterization of *P. hygromyciniae* strain SDM007. **a, b, c.** Optical density (OD_600_) values of SDM007, *P. fluorescens* strain ATCC 17569, and *P. aeruginosa* strain PAO1 in LB medium at **(a**) 4 °C, (**b**) 30 °C, and (**c**) 37 °C over 24 hours. **d, e, f.** CFU in LB medium at (**d**) 4 °C, (**e**) 30 °C, and (**f**) 37 °C over 12 hours. **g.** OD_600_ values of strains grown in LB medium supplemented with 500 μg/mL hygromycin B at 30 °C. **h.** OD_600_ values of SDM007 grown for 24 hours in LB medium containing 0, 100, 500, or 5,000 μg/mL hygromycin B at 30 °C. At 24 h, the differences in CFU were not significant. **i.** Area of maceration of romaine lettuce leaves inoculated with SDM007, ATCC 17569, PAO1, *E. coli* strain TOP10, and PBS after 72 h of incubation at 30 °C. **j, k, l.** Survival of *G. mellonella* larvae when infected with five doses of (**j**) PAO1, (**k**) ATCC 17569, or (**l**) SDM007. **m.** TOP10 CFU recovered after 0 h or 6 h co-incubation with PAO1, ATCC 17569, SDM007, or PBS. **n.** Growth over 24 h of *E. coli* TOP10 expressing the *hph* gene cloned from SDM007. “Vector” refers to the pEX18.AP plasmid (which contains an ampicillin-resistance cassette) with or without the *hph* genes. “in Amp” or “in Hyg” refers to growth in ampicillin or hygromycin, respectively. * = *p*<0.05, ** = *p*<0.01, and *** = *p*<0.001

Because *P. hygromyciniae* was isolated from a vial of hygromycin B, we investigated whether this bacterium used hygromycin B as a carbon or nitrogen source. *P. hygromyciniae* was grown in M9 minimal medium with or without carbon or nitrogen sources and with or without supplementation with hygromycin B. Although *P. hygromyciniae* grew in complete M9 medium supplemented with hygromycin, it was unable to grow in the same medium lacking glucose (carbon) or ammonium chloride (nitrogen), indicating that it cannot use hygromycin B as a carbon or nitrogen source (**Supplemental Fig. 2**). We next examined carbon sources that could be used by *P. hygromyciniae* for growth (**Supplemental Table 1**). Of 190 tested carbon sources, *P. hygromyciniae* showed measurable growth on 74 (39%), including sugars (glucose, fructose, galactose), amino acids (L-alanine, L-arginine, L-histidine), fatty acids (capric acid, propionic acid), and nucleosides (adenosine, uridine). These results indicate that *P. hygromyciniae* is metabolically versatile, with its primary limitation being the requirement for temperatures below 37 °C.

Since several bacteria within the *P. fluorescens* group are associated with the rhizosphere, we investigated whether *P. hygromyciniae* was pathogenic towards plants using a lettuce leaf model^10^. *P. hygromyciniae* was inoculated into the midribs of romaine lettuce leaves. After 72 h, it displayed an area of maceration (AoM) of 6.17 cm^2^, which was smaller than the 11.83 cm^2^ AoM of the known plant pathogen *P. aeruginosa* strain PAO1^11^ but larger than the AoMs of *P. fluorescens* ATCC 17569, *Escherichia coli* TOP10, and PBS (1.83 cm^2^, 1.0 cm^2^, and 0.0 cm^2^, respectively; **Fig. 2i**). These results suggest that *P. hygromyciniae* is a potential plant pathogen.

Since many bacteria in the *P. fluorescens* group are ubiquitous in soil, where they may encounter terrestrial insects, we examined the virulence of *P. hygromyciniae* in a *Galleria mellonella* (greater wax moth) model of infection^12^. When injected into *Galleria* larvae, killing by *P. aeruginosa* PAO1 (**Fig. 2j**) and *P. fluorescens* ATCC 17569 (**Fig. 2k**) were like that of *P. hygromyciniae*. Specifically, *P. hygromyciniae* killed nearly all larvae within 72 h. Even at the lowest inoculum of 1.4 CFU (log 0.15), only a single larva out of 30 survived (**Fig. 2l**). These results indicate that *P. hygromyciniae*, like the other *Pseudomonas* species tested, is highly virulent in the *Galleria* model.

*P. aeruginosa* is a prominent human pathogen, and *P. fluorescens* has caused outbreaks among hospitalized patients^13^. For this reason, we examined the virulence of *P. hygromyciniae* in a mammalian system despite its inability to grow at 37 °C. We incubated this bacterium with A549 lung carcinoma epithelial-like cells and measured cytotoxicity. *P. aeruginosa* PAO1 caused 10% lysis of the cells after 8 h at 37 °C, but *P. hygromyciniae* and *P. fluorescens* ATCC 17569 displayed only ~1% and 0% cell lysis, respectively (**Supplemental Fig. 3**). Given that *P. hygromyciniae* rapidly dies at 37 °C, supernatants from overnight cultures of *P. hygromyciniae* grown at 30 °C were spiked into A549 wells and allowed to incubate for 8 h, but no cell killing was detected above the level of background (**Supplemental Fig. 3**). These data suggest that *P. hygromyciniae* is unlikely to be a human pathogen, as it does not proliferate at normal physiological temperatures, does not kill a representative mammalian cell line, and does not appear to secrete virulence factors capable of eliciting death of this cell line.

Since a rich variety of bacteria inhabit the soil, these bacteria frequently harbor mechanisms to compete against each other. To explore whether *P. hygromyciniae* can successfully compete with other bacteria, a competition assay between *P. hygromyciniae* and *E. coli* was conducted. *P. hygromyciniae* and *E. coli* strain TOP10 were mixed in liquid culture, plated at high density on LB agar plates, and incubated for 6 h at 30 °C. *E. coli* CFU were then enumerated by plating on selective agar. Fewer than half the number of *E. coli* CFU were recovered from the *P. hygromyciniae* plates compared to the *P. aeruginosa* or *P. fluorescens* plates (**Fig. 2m**). These findings indicate that *P. hygromyciniae* has evolved mechanisms that allow it to successfully compete against some other bacterial species.

We next focused on the 250 kb plasmid, designated “pSDM007,” found in *P. hygromyciniae*. Read coverage analysis indicated ~1.4 copies of pSDM007 per genome, suggesting that this is a single-copy plasmid in the majority of cells. A search of the pSDM007 sequence against all Gammaproteobacteria in the NCBI database (taxid: 1236) as of December 29^th^, 2020, yielded a top hit of the *Pseudomonas rhodesiae* strain BS2777 genome (44% coverage and 98.20% identity) and the *P. aeruginosa* plasmid pNK546b (36% query cover and 80.57% identity) (**Supplemental Information 2**). The plasmid contains a number of genes predicted to play a role in conjugation and replication of plasmids (**Supplemental Fig. 4** and **Supplemental Table 2**).

Comparison to the Comprehensive Antibiotic Resistance Database^14^ indicated that pSDM007 contained two genes predicted to confer resistance to aminoglycosides: *hph*, a known hygromycin-B 4-O-kinase (99.68% identity to a known *hph*), and a gene with 72.57% identity to AAC(2’)-IIa, which encodes an aminoglycoside modifying enzyme. To examine whether either were responsible for the hygromycin B resistance exhibited by *P. hygromyciniae*, each of the two genes along with its promoter region was cloned into an isopropylthiogalactoside (IPTG)-inducible expression plasmid. These plasmids were transformed into *E. coli* TOP10 cells, which were then grown in the presence or absence of hygromycin. Resistance to hygromycin B was not conferred by the AAC(2’)-IIa-like gene in either the presence or absence of IPTG (**Supplemental Fig. 5**). However, the *hph* gene did confer growth in the presence of hygromycin B (100 μg/L) even without IPTG induction (**Fig. 2n**). These results indicate that the *hph* gene carried by the pSDM007 plasmid was at least partially responsible for the hygromycin resistance of *P. hygromyciniae*.

Finally, we examined whether *P. hygromyciniae* could be cultured from other commercially available preparations of hygromycin B. A second bottle of hygromycin B was purchased from the same vendor as the original vial. This second vial was from the same lot as the original isolation source. Hygromycin B was also purchased from two other vendors. When unfiltered hygromycin B from each stock was added to individual culture tubes of liquid LB and cultured at room temperature for 72 h, bacterial growth only occurred from the preparation purchased from the original vendor. DNA from a single colony of this growth (designated SDM007_2) was sequenced, and the resulting genome was found to be similar to the original SDM007 genome, differing by ~700 SNVs on the chromosome and by zero SNVs on the pSDM007 plasmid. These findings confirm that the hygromycin vials were indeed the source of *P. hygromyciniae* and that it was not acquired by contamination in our laboratory. They also suggest that a non-clonal population of this bacterium exists at some point in the hygromycin B production chain of this vendor.

In summary, *P. hygromyciniae* is a novel bacterial species related to *P. fluorescens* that was isolated from a factory-contaminated stock of the aminoglycoside hygromycin B. We propose SDM007 as the type strain for this species. Characteristics of SDM007 include its ability to use a variety of compounds as carbon sources, its pathogenicity towards lettuce and *Galleria*, and its ability to inhibit the growth of an *E. coli* strain. SDM007 is unlikely to be a human pathogen due to its inability to survive at 37 °C as well as its lack of cytotoxicity towards a mammalian cell line. This bacterium also carries a large plasmid that contains a gene conferring high-level resistance to hygromycin B, an antibiotic used widely in research laboratories.

## MATERIALS AND METHODS

### Bacterial strains, media, and reagents

*P. hygromyciniae* SDM007 was isolated from a sealed bottle of lyophilized hygromycin B purchased from Sigma-Aldrich Co. (product number H3274-1G, lot number SLBZ3956) in November 2019. A second bottle of hygromycin B of the same product and lot number was purchased from Sigma-Aldrich in February 2020. Other hygromycin B vials were purchased from Dot Scientific, Inc. (product number DSH75020-1, lot number 94491-105646) and Gold Biotechnology (product number H-270-1). Ampicillin was purchased from Acros Organics. *E. coli*, *P. aeruginosa*, *P. fluorescens*, and *P. hygromyciniae* were grown aerobically (250 RPM) in LB broth (10 g/L tryptone, 5 g/L yeast extract, 10 g/L sodium chloride) or on LB agar plates (LB broth supplemented with 16 g/L agar). M9 medium was made as follows: Na_2_HP0_4_ 6.9 g/L, KH_2_PO_4_ 3 g/L, NaCl 0.5 g/L, NH_4_Cl 1 g/L, CaCl_2_ 0.1 mM, MgSO_4_ 2 mM, and glucose 0.5%. A549 (male, human lung epithelial-like) cells were obtained from ATCC and grown at 37 °C in DMEM (Gibco) supplemented with 10 % FBS (GE Healthcare Life Science) under 5 % CO_2_. All strains used in this study are listed in **Supplemental Table 3**.

### Whole-genome sequencing and sequence analysis

Genomic DNA was extracted using a Maxwell 16 Cell DNA Purification Kit (Promega). Genomic DNA libraries were prepared using a Nextera XT kit (Illumina) following the manufacturer’s protocol. Short-read sequencing was performed on a MiSeq platform (Illumina) generating 300 bp paired-end reads. For SDM007 3,348,302 reads were generated totaling 826,593,557 bp with an average read length of 247 bp after adapter trimming. The average fragment size for SDM007 was 804 bp, which includes adapters trimmed from the data while sequencing. Libraries were prepared for long-read sequencing using the ligation sequencing kit (SQK- LSK109, Oxford Nanopore) and were sequenced on the MinION platform using a FLO-MIN106D flow cell (Oxford Nanopore). Nanopore basecalling with Guppy v3.3.3 yielded 27,630 reads that passed quality filtering and totaling 216,452,583 bases with an average read length of 7,834 bp and an L50 value of 5,006. Hybrid de novo assembly of SDM007 was performed by first assembling Nanopore long reads using Flye (v2.5),^15^ which generated two circularized contigs. Next, Illumina short reads were aligned to the contigs using BWA (v0.7.17)^16^ and mismatch and indel errors corrected using Pilon (v1.23)^17^.

Alignment and error correction were repeated until no new corrections were made by Pilon, after a total of five cycles. Prokka v1.12^18^ was used for initial annotations of the SDM007 genome with additional annotations via BLAST^19^ and the Comprehensive Antibiotic Resistance Database^14^ analysis. A phylogenetic tree was constructed using kSNP v3.0.21^20^ based on SNP loci occurring in at least 95% of genomes. Short-read sequencing of SDM007_2 was performed similarly to that of SDM007. The genomes of SDM007, SDM007_2, and plasmid pSDM007 have been deposited in the GenBank database as accession numbers CP070506, JAFFTG000000000, and NZ_CP070507, respectively.

Polymorphisms between the two *P. hygromyciniae* isolates were determined as previously described.^21^ Briefly, Illumina reads from the second isolate were trimmed with Trimmomatic (v0.36)^22^ and aligned to the SDM007 chromosome or plasmid using BWA (v0.7.15).^16^ Variant sites meeting quality criteria and not in repetitive region of the reference sequence were then identified using SAMtools (v0.1.19-44428cd)^23^ and a custom Perl script.

The 16s rRNA gene sequence analysis utilized the NCBI BLAST 16S ribosomal RNA sequence (bacteria and archaea) database^24^. In silico DNA-DNA hybridization utilized the Type Strain Genome Server^6^ DDH *d*_4_ value. All *Pseudomonas* spp. genomes (9,108 as of December 30^th^, 2019) available from the Pseudomonas Genome Database^25^ were downloaded, and a custom script^8^ was used to determine an approximate ANI of each genome with SDM007. A parsimony phylogenetic tree of SDM007 and all genomes with an approximate ANI ?90% was constructed using kSNP v3.0.21^20^ based on SNP loci occurring in at least 95% of the 25 analyzed genomes, with the *P. fluorescens* reference strain SBW25 as the outgroup.

### Growth curves and antimicrobial susceptibility testing

For growth curves, SDM007 bacteria and control bacteria from −80 °C frozen stocks were struck onto LB agar plates with or without 500 μg/mL hygromycin B. Individual colonies were grown overnight in liquid LB medium at 30 °C with shaking and sub-cultured for 3 h in LB. All strains were adjusted to a final OD_600_ of 0.10 and inoculated into 5 mL of culture media. Cultures were incubated at 4 °C, 30 °C, and 37 °C in 250 RPM orbital shakers (Thermo Fisher MaxQ 4000, USA). Samples were taken at set timepoints and diluted as needed. OD_600_ values were obtained by placing 200 μL of each sample into a 96-well plate on a SpectraMax M3 plate reader (Molecular Devices). Bacterial CFUs were assessed by serial dilutions of samples and plating onto LB agar plates for enumeration of colonies after overnight growth at room temperature.

To determine growth rates of strains in the presence of hygromycin B, bacteria were grown as described above except that 0 μg/mL, 100 μg/mL, 500 μg/mL, or 5,000 μg/mL of filter-sterilized hygromycin B was added to the medium. Bacteria were added to these hygromycin-supplemented media to obtain a final OD_600_ of 0.10. A total of 200 μL of the bacterial suspensions were added to a Costar 96-well clear, flat-bottom plate (Corning). Cultures were incubated for up to 48 h at 30 °C with periodic OD_600_ measurements.

Whether SDM007 could utilize hygromycin as a carbon or nitrogen source was tested using standard M9, M9 without ammonium chloride, and M9 without glucose media. Bacterial suspensions in the indicated medium were adjusted to an OD_600_ of 0.10, and 200 μL was added to a Costar 96-well, clear, flat-bottom plate. Cultures were incubated for 24 h at 25 °C with 250 RPM shaking and periodic reading of OD_600_ values using a SpectraMax M3 plate reader.

All growth curves, antibiotic resistance assays, and minimal media assays were performed with at least two technical replicates (per biological replicate) and three biological replicates, and results are shown as means of these data.

### Expression of *hph* and AAC(2’)-IIa-like gene in *E. coli* TOP10

Oligonucleotides were purchased from Integrated DNA Technologies (Iowa, USA). Polymerase chain reactions (PCR) were performed using Phusion High-Fidelity DNA Polymerase (New England Biolabs), and gel-purified products (*hph* and the gene of unknown function with 73% homology to an AAC(2’)-IIa resistance gene) were cloned into the pEX18.AP vector (with ampicillin resistance as a selectable marker) using the Gibson Assembly Cloning Kit (New England Biolabs) per the manufacturer’s instructions. Chemically competent *E. coli* TOP10 cells were transformed with the Gibson construct following manufacturer’s instructions and were plated on LB agar containing 100 μg/mL ampicillin and grown overnight. Plasmids were purified and verified by PCR amplification; the gel-purified products were Sanger sequenced through the Northwestern University NUSeq Sequencing Core for sequence confirmation. TOP10 cells harboring a pEX18.AP empty vector were also transformed using the above protocol without addition of the gel-purified products and used as a control. Verified strains were stored at −80 °C.

Freezer stocks of the strains were grown overnight at 37 °C on LB agar supplemented with 100 μg/mL ampicillin. A single colony of each strain was inoculated into 5 mL LB medium supplemented with 100 μg/mL ampicillin and grown overnight at 37 °C in a 250 RPM orbital shaker. A 500 μL aliquot of the overnight culture was transferred to 5 mL of fresh LB supplemented with 100 μg/mL ampicillin and grown until mid-log phase. The resulting culture was centrifuged for 90 seconds at 21,000 x *g* and the cell pellet was washed in sterile PBS twice, then resuspended in 1 mL of PBS. The resuspended cells were added to 5 mL of LB supplemented with either 100 μg/mL ampicillin or 100 μg/mL hygromycin B for a final OD_600_ of 0.1. Optical densities were monitored over time to determine if the cloned *hph* and AAC(2’)-IIa-like genes provided hygromycin B resistance.

All plasmids and primers used in this study are listed in **Supplemental Table 3** and **Supplemental Table 4**, respectively.

### Lettuce model of infection

Lettuce leaf infection assays were conducted following the protocol of Starkey & Rahme^10^ with minor modifications using store-bought Earthbound Farm’s Organic Whole Romaine Leaves. A 5-μL-aliquot of bacteria adjusted to an OD_600_ of 1.0 was injected into the midrib of the lettuce, and inoculum size was verified by plating and CFU enumeration. Petri dishes containing the inoculated leaves and Whatman no. 1 filter paper soaked in 10 mM MgSO_4_ were individually placed along with a paper towel soaked in 10 mM MgSO_4_ into partially sealed 3.78 L storage bags (Target Corporation). Storage bags containing the petri dishes were then incubated at 30 °C. After 72 h, areas of necrosis were measured. All tests were performed with at least three technical replicates (per biological replicate) and three biological replicates, and results are shown as means of these data.

### Carbon utilization

Growth with different carbon sources was assessed using Phenotypic MicroArray PM1 and PM2 MicroPlates (Biolog) following the manufacturer’s protocol. Optical densities were read using a SpectraMax M3 plate reader after 12 hours of growth. Results are shown as the average of two biological and two technical replicates. OD_410_ values below 0.20 were interpreted to indicate that the carbon source was not utilizable by the bacteria.

### *Galleria mellonella* model of infection

Overnight bacterial cultures were inoculated into fresh LB medium and grown for 1 hour and 45 minutes at 37 °C with shaking. Cultures were pelleted, washed with sterile PBS, and diluted to provide approximately 10, 100, 1,000, 10,000, and 100,000 bacteria in 10 μL inoculums. Bacterial inoculums were verified by plating and CFU enumeration. The 10 μL inoculums were injected into the distal proleg of *G. mellonella* larvae (purchased from Vanderhorst Wholesale, Inc.) weighing between 200 and 350 mg using a LEGATO 100 Syringe Pump (KD Scientific) equipped with an EXEL 29.5-gauge 0.5 mL syringe (Fisher Scientific). Larvae showing immediate discharge of hemolymph or exaggerated movements post-injection were discarded. Injected larvae were placed in petri dishes and monitored at room temperature for up to 100 h. Larvae that did not move in response to external stimuli were scored as dead. Ten *G. mellonella* larvae were infected per condition, and experiments were repeated three times. Cumulative results of all three experiments are shown.

### Bacterial competition assay

Predator strains (*P. hygromyciniae* SDM007, *P. aeruginosa* PAO1, and *P. fluorescens* ATCC 17569) and a prey strain *(E. coli* TOP10) were grown overnight on LB agar plates at 30 °C. A single colony was picked for each strain and inoculated into individual tubes of LB (5 mL), which were grown overnight at 30 °C with shaking (250 RPM). Aliquots of 500 μL were then subcultured into 5 mL of LB and grown at 30 °C with shaking (250 RPM) for 3 h. Cultures were spun down, washed twice with sterile PBS, and adjusted to OD_600_ ~1. A predator and a prey strain were combined at a 5:1 ratio and briefly vortexed. A total of 25 μL of the bacterial mixture was spotted onto LB agar plates, which were incubated for 6 h at 30 °C. The 0 h and 6 h spottings were scraped from the plate, separately resuspended in 100 μL of sterile PBS, and serially diluted onto LB plates with or without 5 μg/mL irgasan (for counter-selection against *E. coli*) for enumeration of colonies. At least two biological replicates each with three technical replications were performed for each experiment.

### Cytotoxicity assay

Cytotoxicity was quantified by measuring lactate dehydrogenase (LDH) release. A549 cells were seeded into 96-well polystyrene tissue culture plates (Corning) in 200 μL DMEM with phenol red and 10% fetal bovine serum. A549 cells were allowed to grow for 22 h, providing approximately 40,000 A549 cells per well. Cells were washed with PBS, and sterile RPMI medium 1640 (Gibco) was added to each well. Bacterial strains were cultured overnight in LB medium with or without hygromycin B (500 μg/mL). Aliquots of the overnight culture were pelleted by centrifugation at 21,000 x *g* for 90 seconds and washed with PBS twice. Bacteria were resuspended in PBS and added to each A459- containing well at a final multiplicity of infection (MOI) of 10. PBS alone was added to some wells to measure background LDH release, and 1% Triton X-100 in PBS to other wells to measure total cell lysis.

The 96-well plate was centrifuged at 500 x *g* for 5 minutes to enhance bacteria-to-cell contact. Infected cells were incubated at 37 °C under 5% CO_2_. After 8 h, the plate was centrifuged at 500 x *g* for 5 minutes to pellet cell debris, and 30 μL of supernatant was removed from the top of each well for measurement of LDH release using a Pierce LDH Cytotoxicity Assay Kit (Thermo Scientific) as per manufacturer’s instructions. Cytotoxicity was calculated as a percentage of total cell lysis (wells with Triton X-100 in PBS) as follows: % cytotoxicity = [(Sample OD_490_ – Sample OD_680_) – (PBS OD_490_ – PBS OD_680_)] / [(Triton X-100 OD_490_ – Triton X-100 OD_680_) – (PBS OD_490_ – PBS OD_680_)] × 100. Tests were performed with at least two biological replicates, each with three technical replicates, and results are shown as means of these data.

### Statistical analysis

*p*-values were calculated from two-tailed student *t*-tests with the help of the Analysis ToolPak addon (Microsoft Excel).

## ACKNOWLEDGEMENTS

ARH reports grants from the National Institutes of Health during the conduct of this study: R01 AI118257, R21 AI153953 K24 AI104831, and R21 AI164254.

---

**Supplemental Figure 1.**
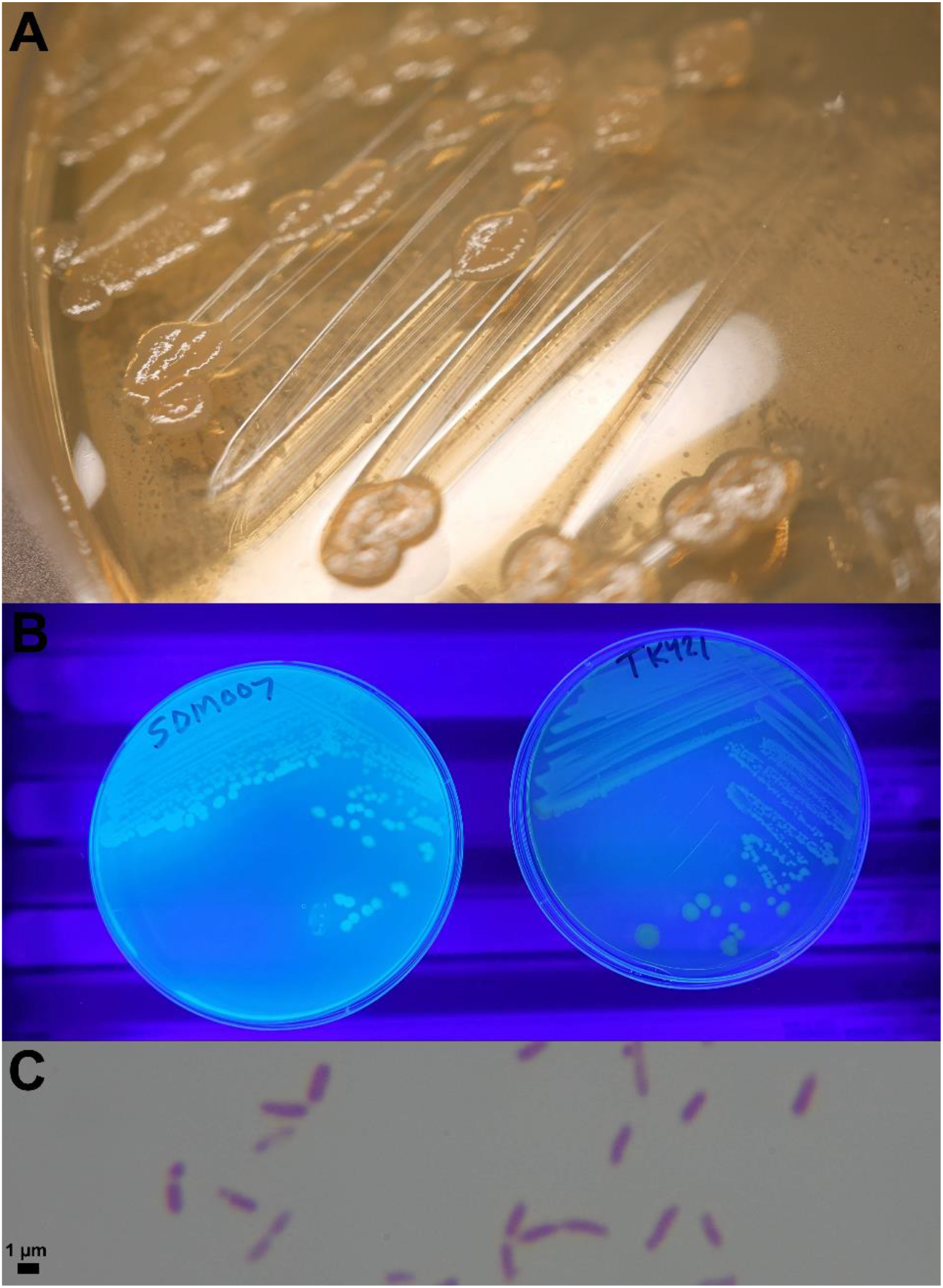
Colony and gram stain characteristics of SDM007. (A) Morphology of SDM007 grown on an LB agar plate. (B) Colony fluorescence of SDM007 (left) and a non-fluorescing control bacterium *Klebsiella pneumoniae* (right) under a UV light. (C) Light microscope image of SDM007 after gram staining, 200x magnification.

**Supplemental Figure 2.**
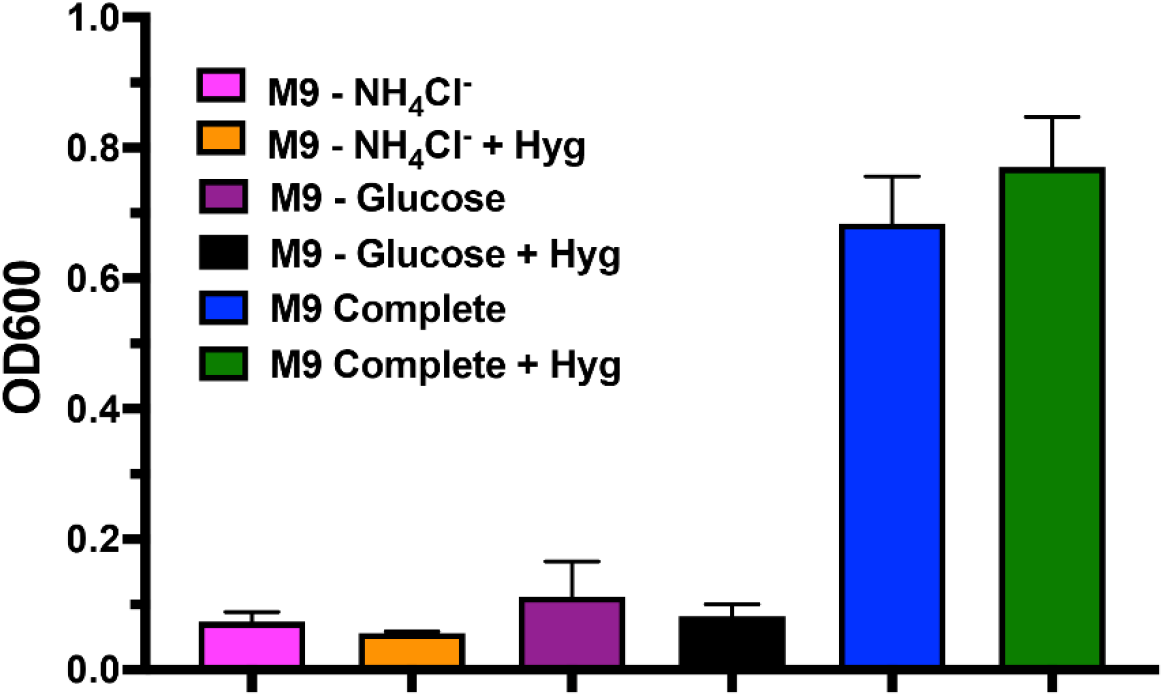
SDM007 growth in minimal medium with or without hygromycin B. M9 minimal medium includes ammonium chloride as a sole nitrogen source and glucose as a sole carbon source. OD_600_ values are shown of SDM007 after growth for 48 h in M9, M9 supplemented with hygromycin B, M9 lacking ammonium chloride, M9 lacking ammonium chloride and supplemented with hygromycin B, M9 lacking glucose, and M9 lacking glucose and supplemented with hygromycin B. Results represent the mean of 2 independent experiments, and error bars indicate standard error.

**Supplemental Figure 3.**
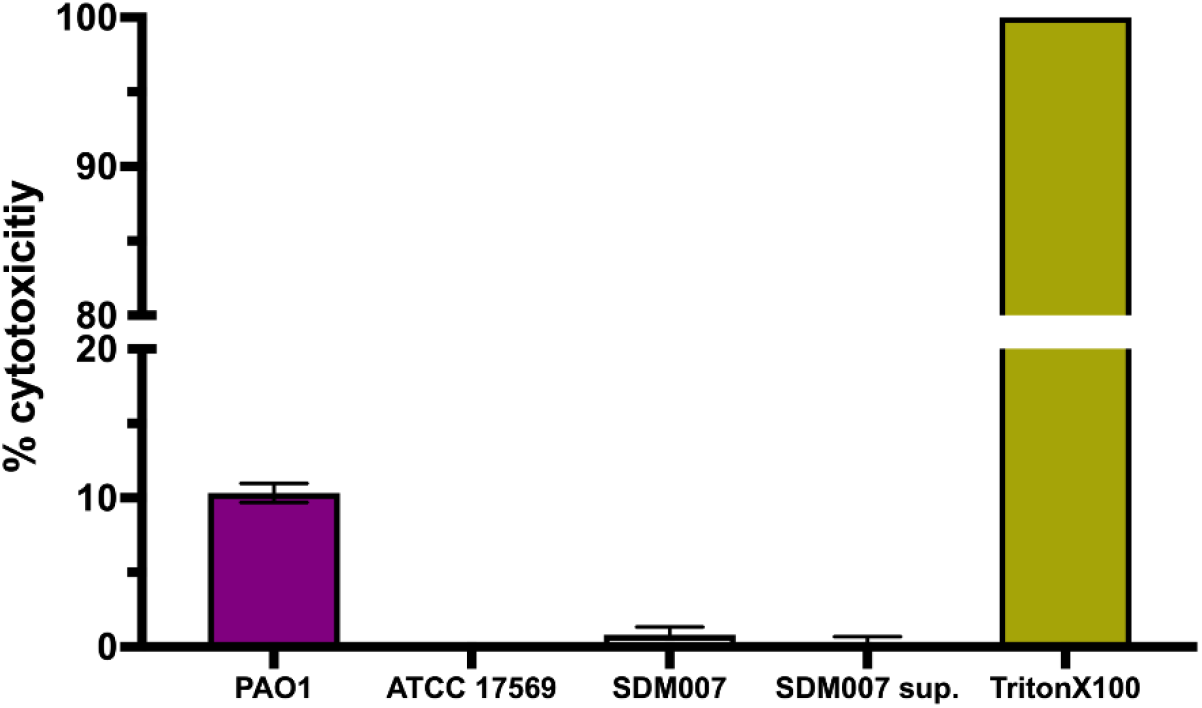
Cytotoxicity of SDM007 towards A549 cells. LDH release from A549 cells incubated for 8 h at 37 °C with *P. aeruginosa* strain PAO1, *P. fluorescens* strain ATCC 17569, SDM007, or the supernatant of an overnight SDM007 culture. Results represent the mean of two independent experiments, and error bars indicate standard error.

**Supplemental Figure 4.**
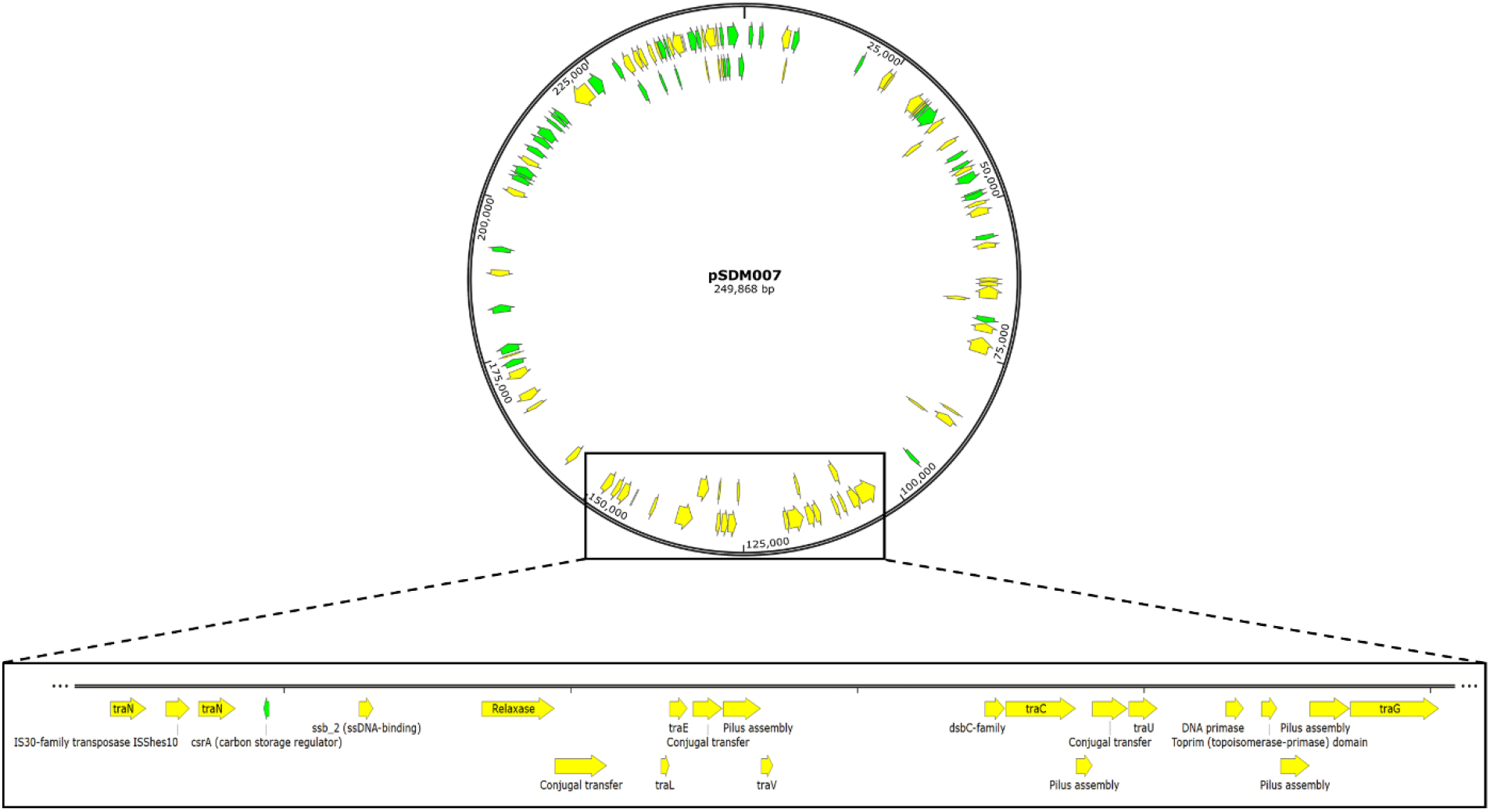
Open reading frames of plasmid pSDM007. The pSDM007 plasmid is 250 Kbp and contains a ~45 Kbp region of putative conjugation genes (see **Supplemental Table 2**).

**Supplemental Figure 5.**
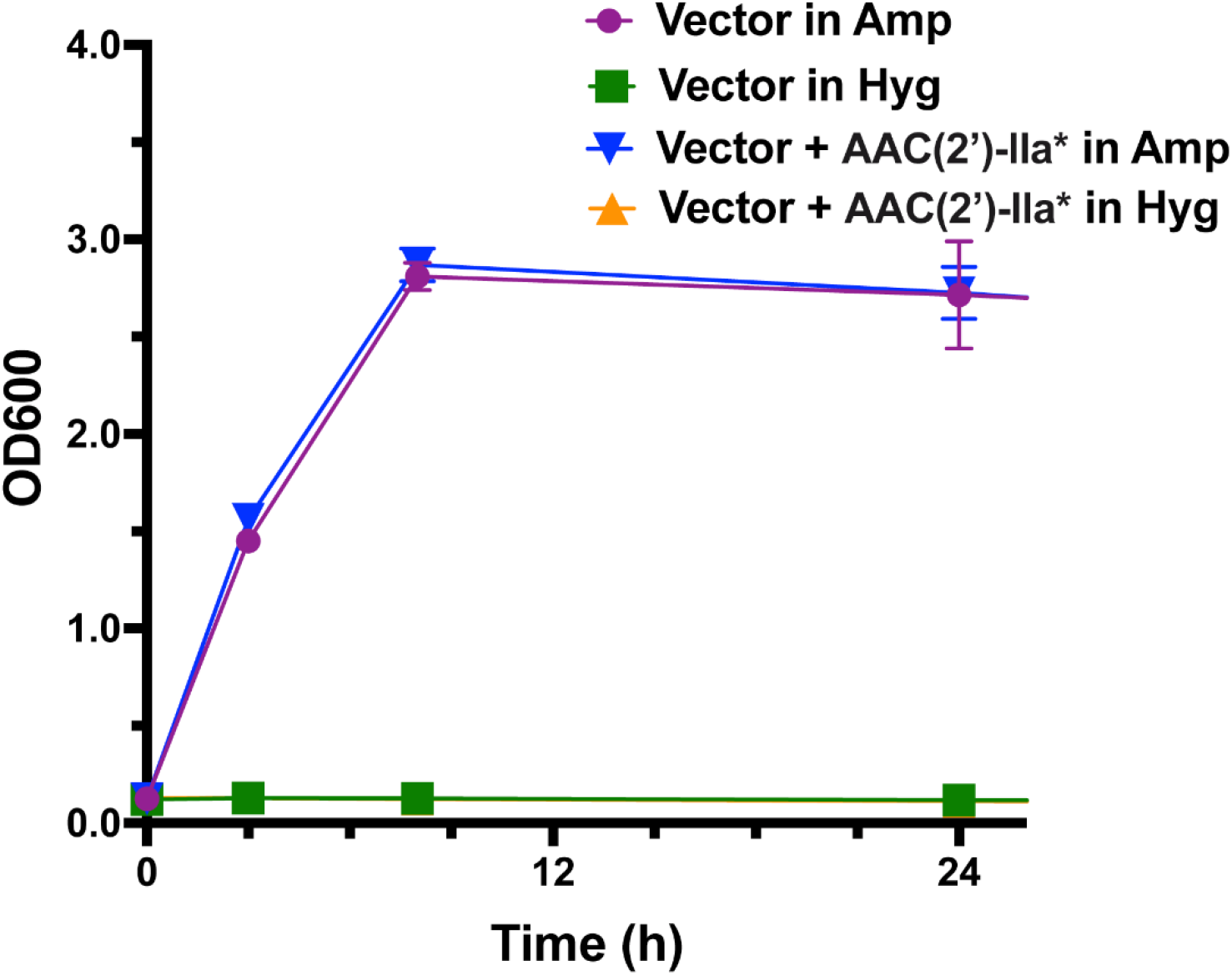
Growth of *E. coli* carrying an AAC(2’)-IIa-like gene from SDM007. *E. coli* TOP10 bacteria were transformed with pEX18.AP (“vector”) containing the AAC(2’)-IIa homolog (“AAC(2’)-IIa*”) or no insert. pEX18.AP expresses an ampicillin selectable marker. The *E. coli* bacteria were then grown in LB medium supplemented with 100 μg/mL hygromycin B (“Hyg”) or 100 μg/mL ampicillin (“Amp”) and OD_600_ values measured over time. Results represent the mean of two independent experiments, and error bars indicate standard error.

**Supplemental Table 1.** Growth of SDM007 on various carbon sources. SDM007 was inoculated into each carbon source using phenotypic microarrays. Growth was measured after 12 hours. Green rows highlight carbon sources that supported growth and red rows highlight caron sources that did not support growth based on a threshold OD_410_ value of 0.200. N = “No or low growth”; Y = “Yes, growth”

| Carbon source | OD<sub>410</sub> value | Standard error | Growth |
| --- | --- | --- | --- |
| 1,2-Propanediol | 0.076 | 0.031 | N |
| 2,3-Butanediol | 0.071 | 0.041 | N |
| 2,3-Butanedione | 0.017 | 0.017 | N |
| 2-Aminoethanol | 0.296 | 0.038 | Y |
| 2-Deoxy adenosine | 0.043 | 0.043 | N |
| 2-Deoxy-D-Ribose | 0.040 | 0.040 | N |
| 2-Hydroxy benzoic acid | 0.008 | 0.008 | N |
| 3-O-β-D-Galactopyranosyl-D-arabinose | 0.057 | 0.033 | N |
| 3-Hydroxy-2-butanone | 0.017 | 0.017 | N |
| 3-Methyl glucose | 0.036 | 0.036 | N |
| 4-Hydroxy benzoic acid | 0.391 | 0.082 | Y |
| 5-Keto-D-gluconic acid | 0.101 | 0.101 | N |
| Acetamide | 0.012 | 0.012 | N |
| Acetic acid | 0.373 | 0.064 | Y |
| Acetoacetic acid | 0.090 | 0.049 | N |
| Adenosine | 0.305 | 0.024 | Y |
| Adonitol | 0.091 | 0.027 | N |
| Amygdalin | 0.020 | 0.020 | N |
| Arbutin | 0.003 | 0.003 | N |
| Bromo succinic acid | 0.472 | 0.062 | Y |
| Butyric acid | 0.012 | 0.012 | N |
| Capric acid | 0.414 | 0.048 | Y |
| Caproic acid | 0.168 | 0.168 | N |
| Chondroitin sulfate C | 0.107 | 0.031 | N |
| Citraconic acid | 0.041 | 0.041 | N |
| Citramalic acid | 0.022 | 0.022 | N |
| Citric acid | 1.053 | 0.001 | Y |
| D,L-Carnitine | 0.376 | 0.013 | Y |
| D,L-Malic acid | 0.686 | 0.030 | Y |
| D,L-Octopamine | 0.020 | 0.020 | N |
| D,L-α-Glycerol-phosphate | 0.427 | 0.002 | Y |
| D-Alanine | 0.743 | 0.039 | Y |
| D-Arabinose | 0.046 | 0.046 | N |
| D-Arabitol | 0.283 | 0.055 | Y |
| D-Aspartic acid | 0.021 | 0.021 | N |
| D-Cellobiose | 0.053 | 0.053 | N |
| Dextrin | 0.059 | 0.043 | N |
| D-Fructose | 0.587 | 0.013 | Y |
| D-Fructose-6-phosphate | 0.056 | 0.042 | N |
| D-Fucose | 0.032 | 0.032 | N |
| D-Galactonic acid-γ-lactone | 0.985 | 0.037 | Y |
| D-Galactose | 0.609 | 0.085 | Y |
| D-Galacturonic acid | 0.035 | 0.035 | N |
| D-Gluconic acid | 1.032 | 0.045 | Y |
| D-Glucosamine | 0.293 | 0.039 | Y |
| D-Glucosaminic acid | 0.077 | 0.040 | N |
| D-Glucose-1-phosphate | 0.028 | 0.028 | N |
| D-Glucose-6-phosphate | 0.010 | 0.010 | N |
| D-Glucuronic acid | 0.021 | 0.021 | N |
| Dihydroxyacetone | 0.423 | 0.040 | Y |
| D-Lactic acid methyl ester | 0.048 | 0.048 | N |
| D-Malic acid | 0.199 | 0.092 | N |
| D-Mannitol | 1.150 | 0.052 | Y |
| D-Mannose | 0.916 | 0.048 | Y |
| D-Melezitose | 0.052 | 0.052 | N |
| D-Melibiose | 0.038 | 0.038 | N |
| D-Psicose | 0.089 | 0.086 | N |
| D-Raffinose | 0.012 | 0.012 | N |
| D-Ribono-1,4-lactone | 1.375 | 0.014 | Y |
| D-Ribose | 0.357 | 0.028 | Y |
| D-Saccharic acid | 0.013 | 0.013 | N |
| D-Serine | 0.930 | 0.001 | Y |
| D-Sorbitol | 0.028 | 0.028 | N |
| D-Tagatose | 0.052 | 0.019 | N |
| D-Tartaric acid | 0.032 | 0.032 | N |
| D-Threonine | 0.048 | 0.048 | N |
| D-Trehalose | 1.043 | 0.040 | Y |
| Dulcitol | 0.032 | 0.032 | N |
| D-Xylose | 0.028 | 0.028 | N |
| Formic acid | 0.077 | 0.036 | N |
| Fumaric acid | 1.033 | 0.026 | Y |
| Gelatin | 0.032 | 0.027 | N |
| Gentiobiose | 0.006 | 0.006 | N |
| Glucuronamide | 0.025 | 0.025 | N |
| Glycerol | 0.702 | 0.069 | Y |
| Glycine | 0.123 | 0.042 | N |
| Glycogen | 0.036 | 0.036 | N |
| Glycolic acid | 0.041 | 0.041 | N |
| Glycolic acid | 0.022 | 0.022 | N |
| Glycyl-L-aspartic acid | 0.034 | 0.034 | N |
| Glycyl-L-glutamic acid | 0.296 | 0.050 | Y |
| Glycyl-L-proline | 0.031 | 0.031 | N |
| Glyoxylic acid | 0.072 | 0.069 | N |
| Hydroxy-L-proline | 1.647 | 0.113 | Y |
| i-Erythritol | 0.254 | 0.009 | Y |
| Inosine | 0.612 | 0.013 | Y |
| Inulin | 0.247 | 0.091 | Y |
| Itaconic acid | 0.000 | 0.000 | N |
| Lactitol | 0.033 | 0.033 | N |
| Lactulose | 0.044 | 0.044 | N |
| L-Alaninamide | 0.909 | 0.022 | Y |
| L-Alanine | 0.751 | 0.027 | Y |
| L-Alanyl-glycine | 0.666 | 0.039 | Y |
| Laminarin | 0.075 | 0.055 | N |
| L-Arabinose | 0.018 | 0.018 | N |
| L-Arabitol | 0.000 | 0.000 | N |
| L-Arginine | 0.509 | 0.060 | Y |
| L-Asparagine | 1.122 | 0.042 | Y |
| L-Aspartic acid | 0.992 | 0.079 | Y |
| L-Fucose | 0.020 | 0.020 | N |
| L-Galactonic acid-γ-lactone | 0.034 | 0.034 | N |
| L-Glucose | 0.013 | 0.013 | N |
| L-Glutamic acid | 0.880 | 0.030 | Y |
| L-Glutamine | 1.068 | 0.054 | Y |
| L-Histidine | 1.127 | 0.055 | Y |
| L-Homoserine | 0.000 | 0.000 | N |
| L-Isoleucine | 1.347 | 0.108 | Y |
| L-Lactic acid | 0.868 | 0.033 | Y |
| L-Leucine | 1.209 | 0.031 | Y |
| L-Lysine | 0.113 | 0.036 | N |
| L-Lyxose | 0.043 | 0.043 | N |
| L-Malic acid | 0.796 | 0.098 | Y |
| L-Methionine | 0.064 | 0.027 | N |
| L-Ornithine | 0.838 | 0.060 | Y |
| L-Phenylalanine | 0.512 | 0.065 | Y |
| L-Proline | 1.007 | 0.055 | Y |
| L-Pyroglutamic acid | 0.769 | 0.035 | Y |
| L-Rhamnose | 0.029 | 0.029 | N |
| L-Serine | 0.888 | 0.070 | Y |
| L-Sorbose | 0.039 | 0.039 | N |
| L-Tartaric acid | 0.045 | 0.023 | N |
| L-Threonine | 0.173 | 0.059 | N |
| L-Valine | 0.347 | 0.017 | Y |
| Malonic acid | 0.450 | 0.006 | Y |
| Maltitol | 0.034 | 0.034 | N |
| Maltose | 0.078 | 0.054 | N |
| Maltotriose | 0.122 | 0.096 | N |
| Mannan | 0.025 | 0.025 | N |
| Melibiononic acid | 0.046 | 0.039 | N |
| Methyl pyruvate | 0.559 | 0.040 | Y |
| m-Hydroxy phenyl acetic acid | 0.032 | 0.032 | N |
| Mono methyl succinate | 0.091 | 0.056 | N |
| m-Tartaric acid | 0.025 | 0.025 | N |
| Mucic acid | 1.510 | 0.143 | Y |
| Myo-inositol | 0.593 | 0.013 | Y |
| N-Acetyl-D-galactosamine | 0.030 | 0.028 | N |
| N-Acetyl-D-glucosamine | 0.566 | 0.060 | Y |
| N-Acetyl-D-glucosaminitol | 0.062 | 0.036 | N |
| N-Acetyl-L-glutamic acid | 1.027 | 0.035 | Y |
| N-Acetyl-Neuraminic acid | 0.009 | 0.009 | N |
| N-Acetyl-β-D-mannosamine | 0.110 | 0.024 | N |
| Negative Control | 0.000 | 0.000 | N |
| Oxalic acid | 0.033 | 0.033 | N |
| Oxalomalic acid | 0.045 | 0.038 | N |
| Palatinose | 0.035 | 0.035 | N |
| Pectin | 0.085 | 0.074 | N |
| Phenylethyl-amine | 0.039 | 0.039 | N |
| p-Hydroxy phenyl acetic acid | 0.650 | 0.049 | Y |
| Propionic acid | 0.607 | 0.011 | Y |
| Putrescine | 0.500 | 0.108 | Y |
| Pyruvic acid | 0.711 | 0.002 | Y |
| Quinic acid | 0.935 | 0.114 | Y |
| Salicin | 0.024 | 0.024 | N |
| Sebacic acid | 0.777 | 0.048 | Y |
| Sec-Butylamine | 0.000 | 0.000 | N |
| Sedoheptulosan | 0.037 | 0.037 | N |
| Sorbic acid | 0.035 | 0.020 | N |
| Stachyose | 0.029 | 0.025 | N |
| Succinamic acid | 1.253 | 0.023 | Y |
| Succinic acid | 0.863 | 0.050 | Y |
| Sucrose | 0.033 | 0.033 | N |
| Thymidine | 0.054 | 0.054 | N |
| Tricarballic acid | 0.028 | 0.028 | N |
| Turanose | 0.044 | 0.035 | N |
| Tween 20 | 0.529 | 0.055 | Y |
| Tween 40 | 0.428 | 0.043 | Y |
| Tween 80 | 0.384 | 0.042 | Y |
| Tyramine | 0.763 | 0.038 | Y |
| Uridine | 0.302 | 0.042 | Y |
| Xylitol | 0.030 | 0.030 | N |
| α-Cyclodextrin | 0.021 | 0.013 | N |
| α-D-Glucose | 1.227 | 0.025 | Y |
| α-D-Lactose | 0.051 | 0.025 | N |
| α-Hydroxy butyric acid | 0.148 | 0.031 | N |
| α-Hydroxy glutaric acid-γ-lactone | 0.655 | 0.061 | Y |
| α-Keto-butyric acid | 0.153 | 0.055 | N |
| α-Keto-glutaric acid | 0.806 | 0.026 | Y |
| α-Keto-valeric acid | 0.066 | 0.066 | N |
| α-Methyl-D-galactoside | 0.068 | 0.047 | N |
| α-Methyl-D-glucoside | 0.040 | 0.022 | N |
| α-Methyl-D-mannoside | 0.055 | 0.055 | N |
| β-Cyclodextrin | 0.031 | 0.031 | N |
| β-D-Allose | 0.005 | 0.005 | N |
| β-Hydroxy butyric acid | 1.142 | 0.091 | Y |
| β-Methyl-D-galactoside | 0.040 | 0.009 | N |
| β-Methyl-D-glucoside | 0.077 | 0.015 | N |
| β-Methyl-D-glucuronic acid | 0.047 | 0.034 | N |
| β-Methyl-D-xyloside | 0.016 | 0.016 | N |
| γ-Amino butyric acid | 1.670 | 0.021 | Y |
| γ-Cyclodextrin | 0.007 | 0.007 | N |
| δ-Amino valeric acid | 0.199 | 0.005 | N |

**Supplemental Table 2.** Putative conjugation genes on plasmid pSDM007. Query coverage, percent identity, and highest similarity were generated using BLAST (blastn) against the NCBI Gammaproteobacteria database. Location (bp) is based on accession number NZ_CP070507 sequence.

| **pSDM007 genes predicted to play a role in conjugation** |  |  |  |  |
| --- | --- | --- | --- | --- |
| **Gene or predicted function** | **Location (bp)** | **Query Cover (%)** | **Identity (%)** | **Highest Similarity** |
| *traG* | 131,019-132,658 | 100 | 84.27 | *P. putida* plasmid pDK1 |
| Pilus assembly | 132,670-134,073 | 99 | 84.96 | *P. rhodesiae* strain BS2777 |
| Pilus assembly | 134,070-135,095 | 100 | 85.38 | *P. aeruginosa* plasmid p1160-VIM |
| DNA primase | 136,378-137,001 | 100 | 88.47 | *P. putida* plasmid pDK1 |
| Nuclease-related domain-containing protein | 138,615-139,334 | 100 | 84.03 | *P. putida* plasmid pDK1 |
| *traU* | 139,384-140,385 | 100 | 86.27 | *P. rhodesiae* strain BS2777 |
| Conjugal transfer protein | 140,420-141,661 | 100 | 82.13 | *P. aeruginosa* plasmid p1160-VIM |
| Pilus assembly protein | 141,655-142,143 | 99 | 85.66 | *P. aeruginosa* plasmid p1160-VIM |
| *traC* DNA transfer | 142,227-144,692 | 100 | 86.54 | *P. aeruginosa* plasmid p1160-VIM |
| *traV* | 152,800-153,225 | 100 | 83.57 | *P. poae* strain CAP-2018 |
| Pilus assembly protein | 153,225-154,546 | 99 | 80.06 | *P. resinovorans* plasmid pCAR1.3 |
| Conjugal transfer protein | 154,565-155,604 | 100 | 80.58 | *P. aeruginosa* plasmid p1160-VIM |
| *traE* | 155,782-156,423 | 100 | 81.93 | *P. aeruginosa* plasmid p1160-VIM |
| *traL* | 145,423-156,710 | 100 | 86.46 | *P. aeruginosa* plasmid p1160-VIM |
| Conjugal transfer protein | 158,590-160,419 | 100 | 76.08 | *P. veronii* strain 1YdBTEX2 |
| Relaxase | 160,416-162,969 | 100 | 82.28 | *P. veronii* strain 1YdBTEX2 |
| *traN* mating pair stabilization | 174,658-175,913 | 100 | 74.78 | *P. aeruginosa* plasmid pNK546b |

**Supplemental Table 3.** Strains and plasmids used in this study

| Species | Strain | Notes | Reference |
| --- | --- | --- | --- |
| *E. coli* | TOP10 | Chemical-competent | Invitrogen |
| *E. coli* | TOP10pEX18.AP | Ampicillin resistant | This study |
| *E. coli* | TOP10pEX18.AP_hph | Ampicillin resistant and harboring SDM007 *hph* gene | This study |
| *E. coli* | TOP10pEX18.AP_ AAC(2')-IIa* | Ampicillin resistant and harboring SDM007 gene of unknown function with 72.57% homology to an AAC(2')-IIa resistance gene | This study |
| *P. aeruginosa* | PAO1 | Canonical *P. aeruginosa* isolate | Holloway<sup>26</sup> |
| *P. fluorescens* | ATCC 17569 | Strain 202 [PJ 372] | ATCC |
| *P. hygromyinae* | SDM007 | Isolated from hygromycin B stock | This study |
| *P. hygromyinae* | SDM007_2 | Isolated from 2<sup>nd</sup> bottle of hygromycin B stock | This study |
| pEX18.AP | n/a | Ampicillin resistance, *oriT*, *sacB*, *lacZα*, MCS from pUC18 | Hoang et al<sup>27</sup> |

**Supplemental Table 4.**
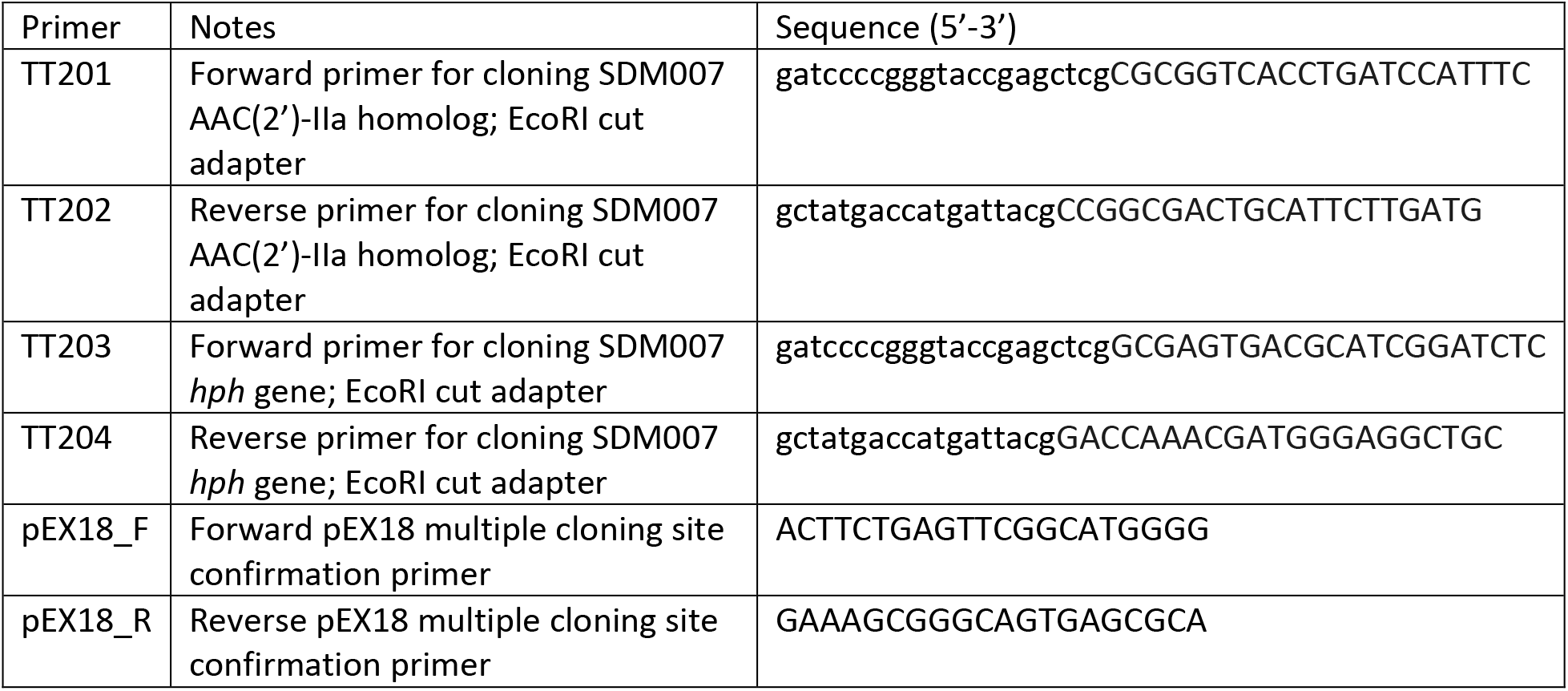
Primers used in this study. For the sequences of primers TT201 – TT204, lowercase letters indicate homology to the pEX18 plasmid, and uppercase letter indicate homology to the AAC(2’)-IIa gene.

